# Identification and Characterization of a Novel Major Facilitator Superfamily (MFS) Efflux Pump SA09310 Mediating Tetracycline Resistance in *Staphylococcus aureus*

**DOI:** 10.1101/2023.01.09.523367

**Authors:** Daiyu Li, Yan Ge, Ning Wang, Yun Shi, Gang Guo, Quanming Zou, Qiang Liu

## Abstract

Drug efflux systems have recently been recognized as an important mechanism of multidrug resistance in bacteria. Here, we described the identification and characterization of a novel chromosomally encoded multidrug efflux pump (SA09310) in *Staphylococcus aureus*. SA09310 is a 43-kDa protein with 12 transmembrane helices. The conserved amino acid sequence motifs of the major facilitator superfamily (MFS) were identified in the protein SA09310, which indicated SA09310 belonged to MFS transporters. Expression of the *sa09310* gene was induced by different types of antibiotics, including aminoglycoside, tetracycline, macrolides, and chloramphenicol. The *sa09310* gene knockout mutant (Δ*sa09310*) was constructed, and its susceptibility to 30 different antibiotics was evaluated. Mutant *△sa09310* exhibited increased sensitivity to tetracycline and doxycycline, with 64-fold and 8-fold decreased MICs, respectively. The mechanism of SA09310 mediating tetracycline resistance was demonstrated by its ability to extrude intracellular tetracycline from within the cells into the environment. The efflux activity of SA09310 was further confirmed by EtBr accumulation and efflux assays. In addition, the efflux activity of SA09310 was observed to be blocked by the known efflux pump inhibitor carbonyl cyanide-chlorophenylhydrazone (CCCP), which provided direct evidence that suggested the H^+^-dependent activity of SA09310 efflux pump. The conservation of SA09310 homologs in Staphylococcus indicated the universal function of these SA09310-like protein clusters. In conclusion, the function-unknown protein SA09310 has been identified and characterized as a tetracycline efflux pump, thereby mediating tetracycline resistance in *S. aureus*.

## Introduction

*Staphylococcus aureus* is a major gram-positive pathogenic bacterium causing a variety of diseases in humans (1). The success of *S. aureus* as a leading pathogen is undoubtedly due to its ability to develop resistance to a wide variety of antimicrobial compounds (2). Antimicrobial resistance of *S. aureus* is mediated by various strategies, including enzymatic modification of the antimicrobial binding site to decrease the affinity of the antibiotic, enzymatic inactivation or degradation of the antimicrobial, decreased permeability of the bacterial cell to antibiotics, and reduction of the intracellular concentration of antibiotics by activating the expression of multidrug efflux pumps to extrude antimicrobial molecules (3-5). Of these, the efflux-mediated resistance has been overshadowed in contrast with the other mechanisms known. However, it has been attracting more interest recently, as it was found that many bacterial efflux pumps were able to recognize and export a broad spectrum of structurally unrelated substrates from the cell, promoting the appearance of multidrug resistance phenotypes (6-9).

Multidrug efflux pumps are membrane-integrated proteins involved in the extrusion of toxic agents, such as antibiotics, biocides, and toxic metals, from within the bacteria into the environment (10). Based on bioenergetic and structural criteria, the multidrug efflux system is classified into five families: the major facilitator superfamily (MFS), the small multidrug resistance (SMR) family, the multidrug and toxic compound extrusion (MATE) family, the resistance-nodulation-cell division (RND) superfamily, and the adenosine-triphosphate (ATP)-binding cassette (ABC) superfamily (11). The transporters of the first four families are secondary transporters that use an electrochemical gradient, typically proton motive force, as the driving force for transport, while the transporters of the ABC family are the primary transporters that use ATP to drive the extrusion of their substrates (12). Within these multidrug efflux systems, the MFS has been the most extensively studied among staphylococcal multidrug efflux pumps, which include NorA, NorB, NorC, Tet38, LmrS, SdrM, and MdeA (10). Staphylococcal MFS transporters typically contain 380-480 amino acids that are arranged into 12 or 14 transmembrane segments (TMS), with a conserved MFS-specific motif that lies in the cytoplasmic loop between the TMS2 and TMS3 helices (13). In addition to MFS transporters, SMR transporters including QacD, QacG, QacH, and QacJ, are encoded on the plasmid; ABC transporters including AbcA, Sav1866, and MepA transporters belonging to the MATE family were characterized as being involved in antibiotic resistance in *S. aureus* (10).

Based on the in silico analysis, the *S. aureus* chromosome encoded 31 multidrug efflux pumps that covered all five families (14). However, only one-third (10/31) of them have been studied previously and most of these efflux pumps are function unknown (14). Efflux-mediated multidrug resistance, particularly in staphylococci, is an urgent clinical problem, rendering many of the current antimicrobials ineffective. Thus, inhibition of bacterial multidrug efflux pumps is a reasonable strategy to combat multidrug-resistant *S. aureus*. This potential strategy promoted the study of the identification and development of efflux pump inhibitors for *S. aureus* (15, 16). Therefore, a more in-depth understanding of efflux pump function, regulatory mechanisms, and association with clinical antibiotic resistance will enable the design of better antibiotics that will be less susceptible to bacterial resistance.

In this report, we described the identification and characterization of the *sa09310* gene coding protein with 12 predicted transmembrane helices belonging to the MFS transporter group. We hypothesized that SA09310 is a multidrug efflux pump and investigated its role in mediating antibiotic resistance in *S. aureus*.

## Results

### Gene *sa09310* encodes a transmembrane protein belonging to the MFS transporter family

In the genome of *S. aureus* USA300_FPR3757, the gene *sa09310* (gene locus: SAUSA300_09310) with a length of 1182 bp was predicted to encode a transmembrane protein with a molecular mass of 43.3 kDa. Similar to most MFS transporters (17), the *sa09310* gene coding protein (SA09310) exhibited 12 transmembrane segments based on the prediction by TMHMM 2.0 (Fig. 1A). MFS transporters are characterized by a highly conserved amino acid sequence motif, called motif A, consisting of residues “G(X)_3_DK/RXGRR/K” that lies in the cytoplasmic loop between the TMS2 and TMS3 helices (13, 18, 19). As shown by the transmembrane topology of SA09310, the motif A was also found between TMS2 and TMS3 of SA09310 (Fig. 1A). In addition, another well-conserved motif C, “G(X)_8_ G(X)_3_GP(X)_2_GG”, harbored by drug-ion antiporters of MFS within their fifth α-helix (TMS5)(19, 20), was identified in the fifth TMS of SA09310 as well (Fig. 1A). All this evidence indicated that protein SA09310 was an MFS transporter.

**FIG 1.**
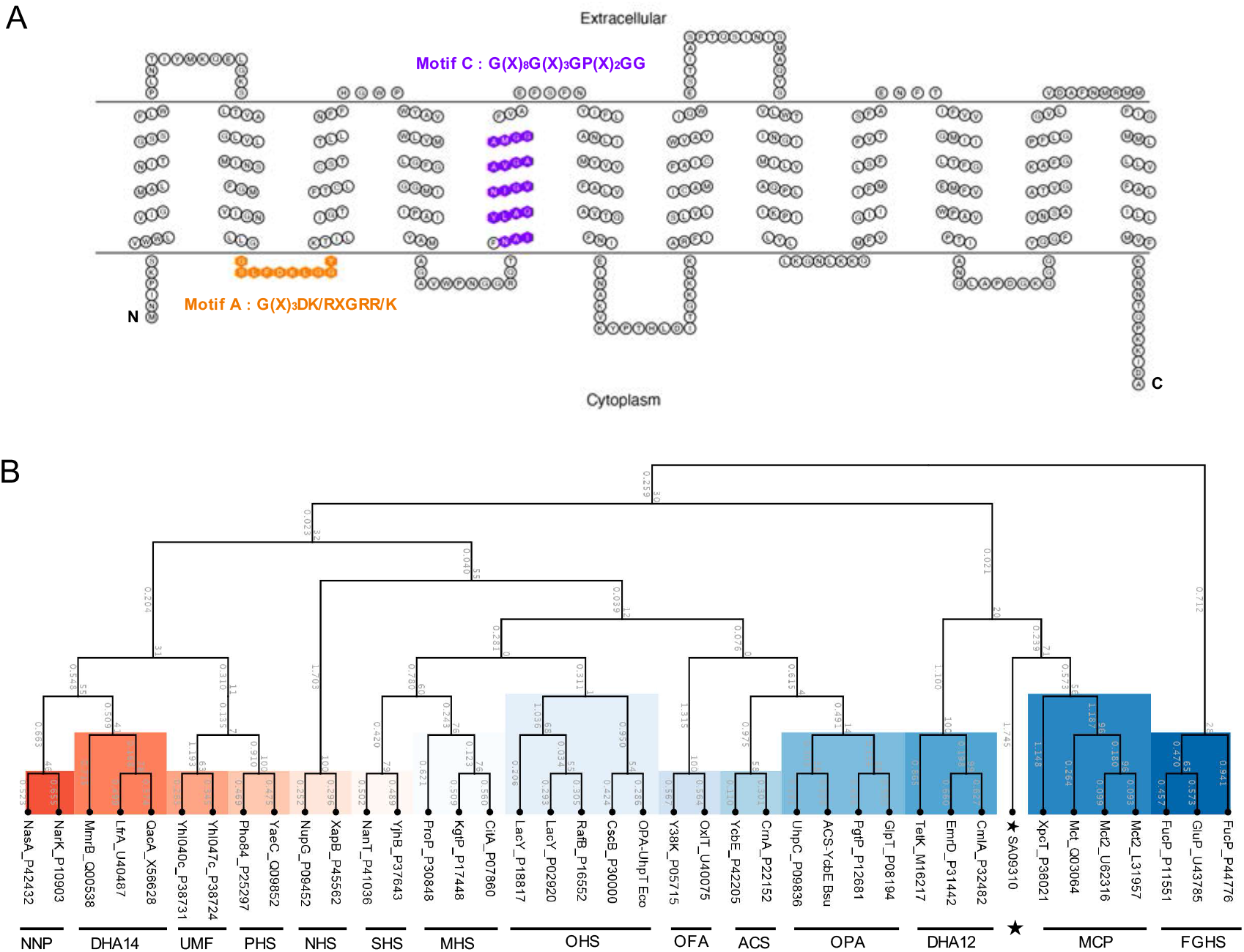
Transmembrane topology and phylogenetic analysis of the SA09310 protein. (A) The transmembrane topology of the SA09310 protein was created using the TOPO2 online tool based on the predicted transmembrane structure by TMHMM 2.0. The amino acids of SA09310 highlighted by orange color indicate the conserved motif (motif A) of MFS transporters between the TMS2 and TMS3 helices. The amino acids of SA09310 highlighted in purple color indicated another conserved motif of MFS transporters located in the fifth α-helix. (B) The phylogenetic tree of SA09310 was constructed based on the protein sequence alignment between SA09310 and proteins from 14 groups of classified MFS transporters using ClustalW, as implemented in the CLC Main Workbench. The resulting tree was calculated using the neighbor-joining method and Jukes-Cantor protein distance model. The abbreviated name of each classified group was indicated in the materials and methods section, and SA09310 was highlighted with a star.

Currently, the MFS transporters are classified into 17 distinct groups based on their protein sequence similarity and distinct functions (17). An unrooted phylogenetic tree was constructed based on the protein sequence alignment between SA09310 and the classified MFS transporters. SA09310 exhibited a close relationship to the drug-H^+^ antiporter (12-spanner) drug efflux (DHA12) and monocarboxylate porter (MCP) families (Fig. 1A), which provided us further clues for determining the function of SA09310.

### Induction of the *sa09310* gene by different types of antibiotics

Since SA09310 was predicted to be an MFS transporter that is phylogenetically close to the drug efflux pump, the role of the *sa09310* gene in response to different antibiotics in *S. aureus* was initially investigated. *S. aureus* cells from the log phase were treated with different types of antibiotics, and the transcriptional level of the *sa09310* gene was analyzed by RT-qPCR. Among the tested antibiotics, quinolone (norfloxacin), glycopeptide (vancomycin), and beta-lactam (ampicillin, oxacillin) were unable to induce the expression of *sa09310* (Fig. 2A). However, *sa09310* was significantly induced by treatment with aminoglycoside (gentamicin, kanamycin), tetracycline, macrolide (erythromycin), and chloramphenicol antibiotics (Fig. 2A). Tetracycline stimulated the *sa09310* gene with the highest (10-fold) transcription level (Fig. 2A). In addition, transcription of *sa0931* induced by tetracycline occurred in a concentration-dependent manner (Fig. 2B), which further confirmed that *sa09310* was able to respond to the stimulation of tetracycline. The response of *sa09310* to different antibiotics suggested that the SA09310 efflux pump might be involved in the extrusion of these antibiotics, which probably mediates the associated antibiotic resistance.

**FIG 2.**
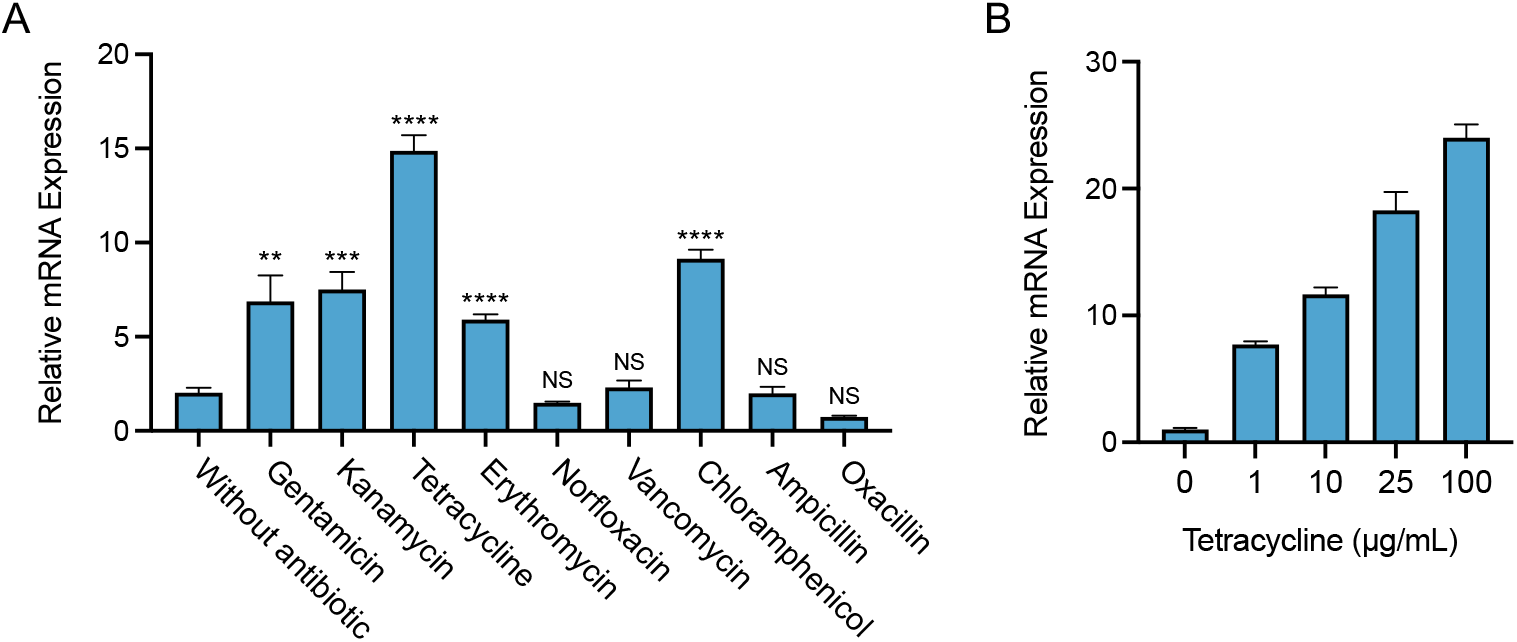
Expression of the *sa09310* gene with the stimulation of different types of antibiotics. (A) Bacterial cells from the mid-exponential phase were treated with 100 µg/mL of different antibiotics. After induction for 1 h, the mRNA levels of the *sa09310* gene were determined by RT-qPCR with *16S rRNA* gene as the internal control. (**** P⩽0.0001; *** P⩽0.001; ** P⩽0.05 relative to the level without antibiotic induction; NS, not significant). (B) The expression of the *sa09310* gene was induced by tetracycline in a concentration-dependent manner. *S. aureus* cells were induced with different concentrations of tetracycline (0, 1, 10, 25, or 100 µg/mL), and the mRNA levels of the *sa09310* gene were checked by RT-qPCR.

### SA09310 contributes to tetracycline resistance in *S. aureus*

Considering the *sa09310* gene was induced by different types of antibiotics, the role of *sa09310* in the antimicrobial resistance of *S. aureus* was subsequently investigated. The *sa09310* gene knockout mutant (Δ*sa09310*) and the strain that complemented *sa09310* gene in Δ*sa09310* (Δ*sa09310*_com) were generated. The susceptibilities of *S. aureus* USA300 WT, Δ*sa09310*, and Δ*sa09310*_com against 32 different antibiotics were tested by disk diffusion assay. The diameter of the inhibitory zone of most antibiotics presented no difference among the *S. aureus* WT, Δ*sa09310*, and Δ*sa09310*_com strains (Table 1), even including gentamicin, kanamycin, erythromycin, and chloramphenicol, which significantly induced the expression of the *sa09310* gene. However, the Δ*sa09310* mutant exhibited significantly larger inhibitory zones than the WT when treated with the disks containing tetracycline (27 ± 0.5 mm for Δ*sa09310* and 9.5 ± 0.5 mm for WT) or doxycycline (27 ± 0.5 mm for Δ*sa09310* and 14 ± 0.5 mm for WT) (Table 1). This phenotype of an enlarged inhibitory zone on Δ*sa09310* plate was able to be restored by complementary expression of the *sa09310* gene in Δ*sa09310* (Δ*sa09310*_com) (Table 1). These results suggested that the *sa09310* gene was involved in tetracycline and doxycycline resistance in *S. aureus*.

**Table 1.** Antimicrobial susceptibility of *S. aureus* WT and its mutant strains in disk diffusion assay

| Antimicrobial agent | Zone diameter of inhibition (mm<sup>a</sup>) |  |  |
| --- | --- | --- | --- |
| | WT | $\Delta sa09310$ | $\Delta sa09310\_com$ |
| Penicillin G (10 U) | 24 ± 0.5 | 24.5 ± 1 | 23.5 ± 0.5 |
| Oxacillin (1 µg) | 17 ± 0.5 | 17.5 ± 0.5 | 17 ± 0.5 |
| Ampicillin (10 µg) | 19.5 ± 0.5 | 20 ± 0.5 | 21 ± 0.5 |
| Carbenicillin (100) | 26 ± 0.5 | 27 ± 0.5 | 27.5 ± 0.5 |
| Piperacillin (100 µg) | 21.5 ± 0 | 22.5 ± 0.5 | 22.5 ± 0.5 |
| Cephalexin (30 µg) | 15 ± 0.5 | 17 ± 0 | 16.5 ± 0 |
| Cefazolin (30 µg) | 24 ± 0.5 | 23.5 ± 1 | 24 ± 0.5 |
| Cefradine (30 µg) | 19 ± 0 | 18.5 ± 0.5 | 18 ± 0.5 |
| Cefuroxime (30 µg) | 18 ± 0.5 | 19 ± 0 | 18 ± 0 |
| Ceftazidime (30 µg) | 13 ± 0.5 | 14 ± 1 | 13 ± 0.5 |
| Ceftriaxone (30 µg) | 16 ± 0.5 | 17.5 ± 0.5 | 16.5 ± 0.5 |
| Cefoperazone (75 µg) | 18.5 ± 0 | 19 ± 0 | 18 ± 0 |
| Amikacin (30 µg) | 14.5 ± 0.5 | 15.5 ± 0 | 15.5 ± 0 |
| Gentamicin (10 µg) | 15.5 ± 0.5 | 16 ± 0 | 16 ± 0 |
| Kanamycin (30 µg) | 15.5 ± 0.5 | 16.5 ± 0.5 | 16 ± 0 |
| Neomycin (30 µg) | 15 ± 0.5 | 16 ± 1 | 15.5 ± 0.5 |
| Tetracycline (30 µg) | 9.5 ± 0.5 | 27 ± 0.5 | 10 ± 0.5 |
| Doxycycline (30 µg) | 14 ± 0.5 | 27 ± 0.5 | 14 ± 0 |
| Minocycline (30 µg) | 24 ± 1 | 25.5 ± 0.5 | 24 ± 0.5 |
| Erythromycin (15 µg) | N | N | N |
| Midecamycin (30 µg) | N | N | N |
| Norfloxacin (10 µg) | N | N | N |
| Ofloxacin (5 µg) | 15.5 ± 0.5 | 16.5 ± 1 | 15 ± 0 |
| Ciprofloxacin (5 µg) | 12 ± 0.5 | 11.5 ± 1 | 10.5 ± 0.5 |
| Vancomycin (30 µg) | 16 ± 0.5 | 17 ± 0.5 | 16 ± 0.5 |
| Polymyxin B (300 U) | N | N | N |
| Sulfamethoxazole (30 µg) | 21 ± 0 | 21 ± 0.5 | 21 ± 0 |
| Furazolidone (300 µg) | 17 ± 0 | 16.5 ± 0.5 | 17.5 ± 0 |
| Chloramphenicol (30 µg) | 21 ± 1 | 19 ± 1 | 19 ± 1 |
| Clindamycin (2 µg) | N | N | N |
| Streptomycin (1 mg) | 22.5 ± 0.5 | 21.5 ± 1 | 22 ± 0.5 |
| Apramycin (0.5 mg) | 21 ± 0.5 | 21 ± 0.5 | 20.5 ± 1 |

<sup>a</sup> Sensitivities were assessed by measuring the diameters of the zones of growth inhibition in three independent assays, and the means ±1 SD were calculated.

Next, the MICs of tetracycline and doxycycline to *S. aureus* WT, Δ*sa09310*, and Δ*sa09310*_com were tested and compared by using E-test method. As shown, strain Δ*sa09310* displayed a tetracycline MIC of 0.125 µg/mL, which decreased 64-fold compared to the WT strain with a MIC of 8 µg/mL (Fig. 3). The same phenotype was observed for the doxycycline strips as well, with the MIC of 0.125 µg/mL for Δ*sa09310* mutant and 1 µg/mL for WT (Fig. 3). Decreased MICs of tetracycline and doxycycline in Δ*sa09310* were able to be restored to WT as indicated by Δ*sa09310*_com as well (Fig. 3). Taken together, these results confirmed that the *sa09310* gene is involved in tetracycline and doxycycline resistance in *S. aureus*.

**FIG 3.**
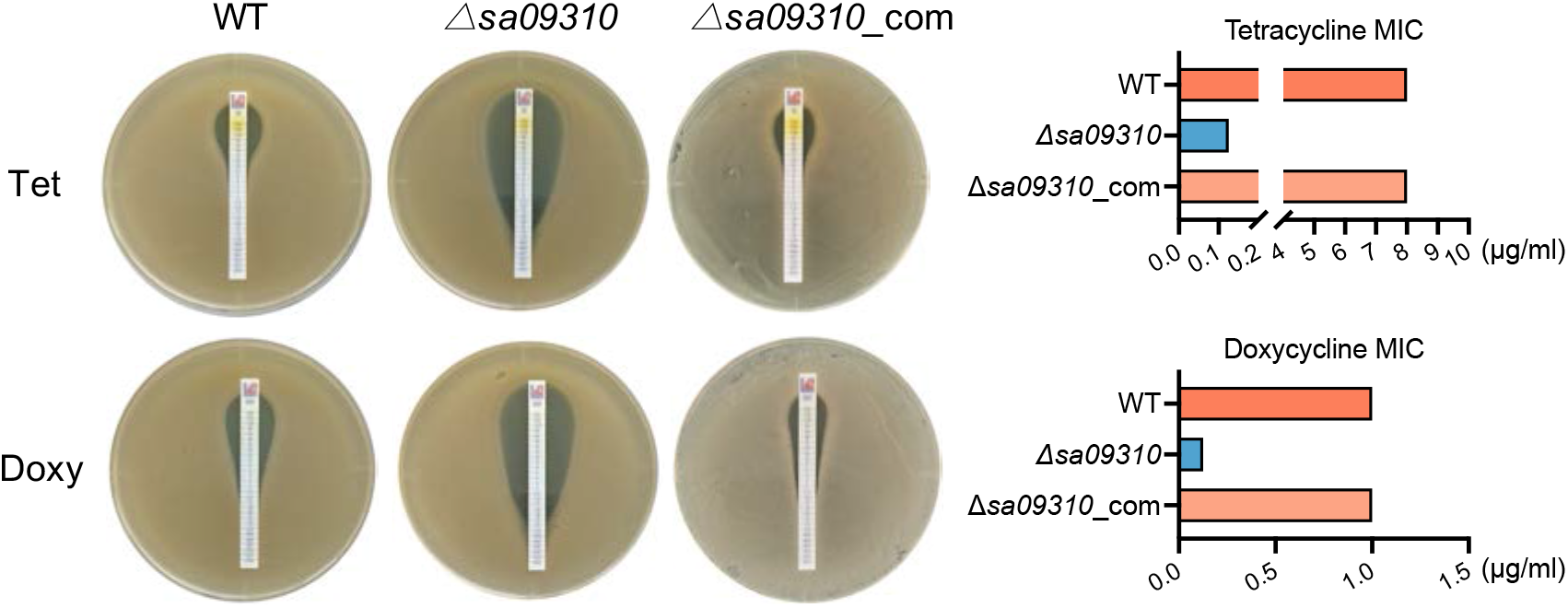
Tetracycline and doxycycline MICs assay. MICs of tetracycline (Tet) and doxycycline (Doxy) were tested with *S. aureus* WT, *sa09310* gene knockout mutant (*△sa09310*), and *△sa09310* complementary (*△sa09310_*com) strains using the E-test Strip on TSA plates according to CLSI guidelines. The MICs of Tet and Doxy for each strain were readed from the plate and presented as histograms in the right panel.

### The contribution of SA09310 to tetracycline resistance is efflux-mediated

Knockout of the *sa09310* gene conferred *S. aureus* with increased sensitivity to tetracycline, which is known to exert antimicrobial activity by targeting the intracellular ribosome (21-23). We hypothesized that SA09310 might play a role as an efflux pump that extruded tetracycline from within the bacterial cells into the environment. To verify this hypothesis, *S. aureus* WT and the Δ*sa09310* mutant were treated with 5 µg/ml tetracycline, and the kinetics of the tetracycline concentration within the bacterial cells of each strain were quantified by using ELISA. As a result, the intracellular tetracycline concentrations in the Δ*sa09310* mutant were 2-to 3-fold higher than those in the WT (Fig. 4A). This result directly supported that SA09310 was an efflux pump that mediated tetracycline resistance of *S. aureus* by extruding intracellular tetracycline.

**FIG 4.**
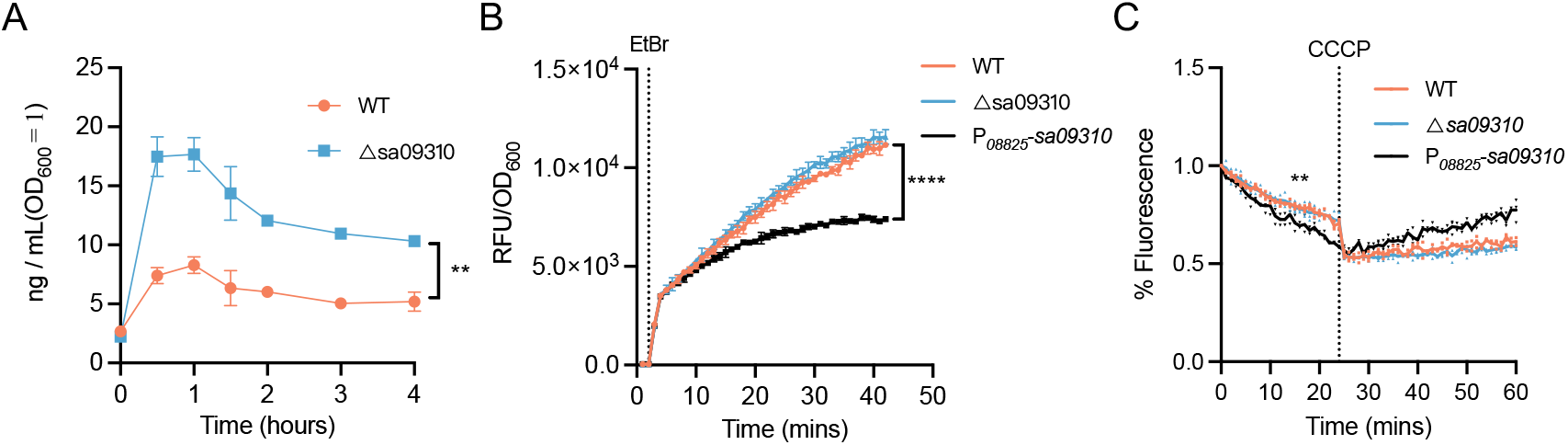
Intracellular tetracycline, EtBr accumulation, and efflux assays. (A) *S. aureus* WT and the Δ*sa09310* mutant were treated with a final concentration of 5 µg/mL tetracycline. At the indicated time points (0, 0.5, 1, 1.5, 2, 3, and 4 h) after treatment, the bacterial cells were harvested and resuspended in lysis buffer by adjusting the OD_600_ value to 1.0 (5×10^8^ CFU/mL). The tetracycline concentration of the cell lysate was quantified by ELISA. Concentrations from suspensions of WT and Δ*sa09310* groups during the indicated period were compared and analyzed. (** P⩽0.05 by paired T-test). (B) EtBr accumulation assay. *S. aureus* WT, Δ*sa09310*, and *sa09310* overexpression strains were treated with 4 µg/mL EtBr. The fluorescence of each strain was measured in a 96-well plate by the Bioreader and normalized to the OD_600_. The dotted line indicates the time point at which EtBr was added. Data are presented as the mean of three independent assays. (**** P⩽0.0001 by paired T-test). (C) EtBr accumulation efflux assay. *S. aureus* cells were treated with EtBr as described above for 40 mins. Then, the extracellular EtBr was removed by centrifugation and resuspension of the cells in fresh PBS, and the fluorescence of each strain was measured. After 20 mins, the efflux pump inhibitor carbonyl cyanide-chlorophenylhydrazone (CCCP) at a final concentration of 100 µM was added to each well, and the fluorescence was read for an additional 40 mins. The dotted line indicated the time point when CCCP was added. (** P⩽0.05 by paired T-test). All data are presented as the mean of three independent assays.

To further confirm the efflux activity of SA09310, EtBr accumulation and efflux assays were performed on WT, Δ*sa09310*, and *sa09310* overexpression strains. In the EtBr accumulation assay, WT, Δ*sa09310*, and *sa09310* overexpression strains were exposed to EtBr, and the fluorescence of each strain was monitored. As shown, a time-dependent increase in fluorescence was observed for all strains, with no difference between WT and Δ*sa09310* (Fig. 4B). However, the *sa09310* overexpression strain displayed a significantly lower increase in fluorescence when compared to WT and Δ*sa09310* (Fig. 4B), which suggested a stronger efflux activity of *sa09310* overexpression strain compared to WT and Δ*sa09310*.

In the EtBr efflux assay, the *sa09310* overexpression strain showed a slightly lower fluorescence compared to WT and Δ*sa09310* (Fig. 4C). This result indicated that overexpression of *sa09310* gene enhanced the efflux activity of *S. aureus*. After the addition of the protonophore CCCP, which dissipates the electrochemical potential of H^+^ across the cytoplasmic membrane, a driving force for the MFS transporters, the fluorescence of all tested strains stopped decreasing due to the collapse of the proton gradient across the membrane (Fig. 4C). This observation provided evidence that suggested H^+^-dependent activity of SA09310. Taken together, SA09310 is proved to be an active efflux pump that mediates tetracycline resistance in *S. aureus* via extrusion of intracellular tetracycline.

### Deletion of *sa09310* gene promote clearance of *S. aureus* by tetracycline in a *G. mellonella* infection model

Knockout of the *sa09310* gene rendered *S. aureus* more sensitive to tetracycline in an *in vitro* assay. However, whether the SA09310 efflux pump promotes tetracycline resistance *in vivo* is requires further investigation, as this information will provide direct evidence of its potential as a target for developing an efflux inhibitor for clinical use. Thus, the clearance of *S. aureus* WT or the Δ*sa09310* mutant by tetracycline in a *G. mellonella* larvae infection model was evaluated.

To determine the optimal infection doses for the *S. aureus* clearance assay, *G. mellonella* larvae were infected with different doses of *S. aureus*. As a result, mortality at 2 days was 100% at 10^7^ CFU/larva and was between 60% and 80% at 10^6^ CFU/larvae, while the group infected with 10^5^ CFU/larva resulted in 90% survival (Fig. 5A). Therefore, the maximum infection doses of 10^5^ CFU/larva that resulted in a survival rate above 80% was used in the clearance assay.

**FIG 5.**
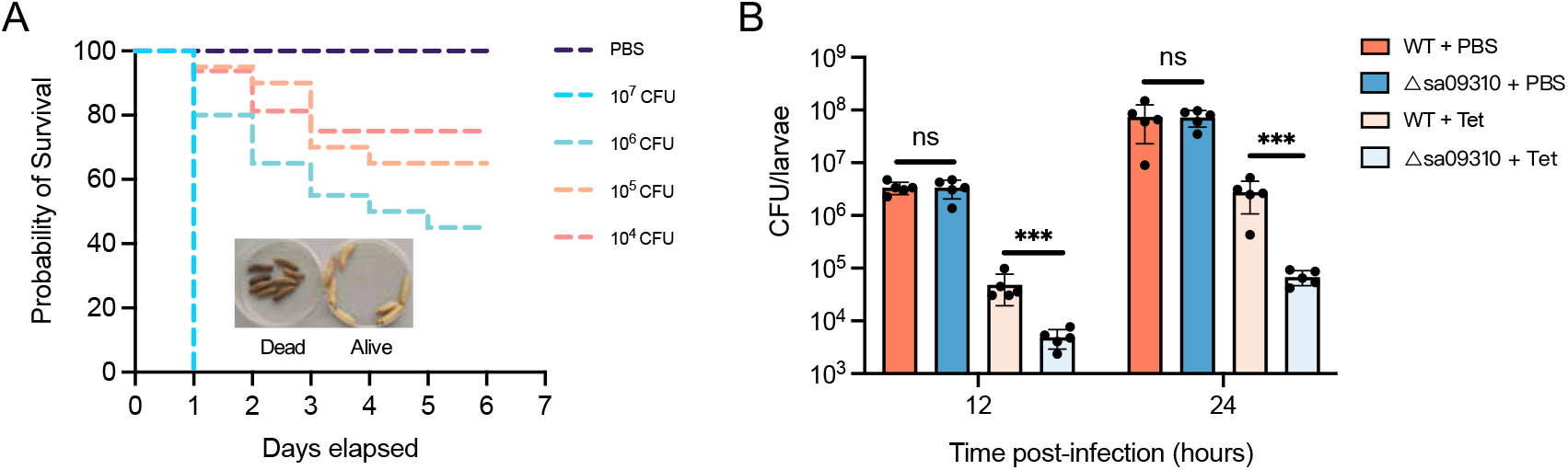
(A) Survival curves of *G. mellonella* larvae infected with *S. aureus* USA300 WT at the dose ranging from 1×10^4^ to 1×10^7^ per larva. Larvae were incubated at 37°C, and their viability was assessed over 6 days. (B) Clearance of *S. aureus* WT and Δ*sa09310* mutant by tetracycline in the larval infection model. Larvae were infected with an optimal dose of 1×10^5^ CFU/larva WT or Δ*sa09310*. After 2 h of infection, 10 µg tetracycline was injected into each larva (tetracycline-treated group). PBS was administered as the control for antibiotic treatment. Five live larvae were randomly selected from each group and were homogenized. The bacterial burden of each selected larva at 12 and 24 h after infection was determined by serial dilution and plating assays. (Significant differences were defined as *** P⩽0.001, ns, not significant)

The efficiency of *S. aureus* WT or the Δ*sa09310* mutant cleared by tetracycline in larvae was evaluated by injecting the larvae with tetracycline after 2 h of infection. First, we confirmed that colonization of *S. aureus* WT and Δ*sa09310* strains exhibited no difference in larvae, as larvae from WT-or Δ*sa09310*-infected group possessed the same bacterial burdens after 12 or 24 h of infection (Fig. 5B). When comparing the infected larvae treated with tetracycline or PBS, the bacterial burdens in the tetracycline-treated groups were significantly lower than PBS-treated groups in both the WT and Δ*sa09310* infection groups (Fig. 5B), which suggested that tetracycline was able to eliminate *S. aureus* efficiently in larvae infection model.

Notably, when comparing the WT- and Δ*sa09310*-infected groups followed by tetracycline treatment, the bacterial burden in the Δ*sa09310*-infected group was 10-to 40-fold lower than that in the WT-infected group (Fig. 5B). As shown, the WT-infected group possessed bacterial burdens of 4.83 ×10^4^ CFU/larva while Δ*sa09310*-infected group carried 4.86 ×10^3^ CFU/larva after 12 h of infection (Fig. 5B). This difference was observed after 24 h of infection as well, in which the WT-infected group had the bacterial burdens of 2.77 ×10^6^ CFU/larva and the Δ*sa09310*-infected group possessed 6.8 ×10^4^ CFU/larva (Fig. 5B). All these results suggested that deletion of the *sa09310* gene renders *S. aureus* significantly prone to be cleared by tetracycline *in vivo*.

### Conservation of SA09310-like efflux pump in staphylococci

To explore the universality of the SA09310 efflux pump in staphylococci, the protein sequence of SA09310 was submitted to the STRING database to identify its homologs among the Staphylococcus genus. As a result, the homologs of SA09310 were identified in all 29 Staphylococcus species. The genomic context of these efflux pump-encoding genes presented a conserved pattern, which included genes encoding a Rot/MarR transcription regulator, methyltransferase domain-containing protein, and TIGR01212 family radical SAM protein upstream of *sa09310*, and a leucine-tRNA ligase-encoding gene downstream of *sa09310* (Fig. 6A). Moreover, the protein sequence of SA09310 from *S. aureus* displayed a 64.6% to 79.6% identity to its analogs from other Staphylococcus species (Fig. 6B). These observations implied that SA09310 and its analogs potentially function the same as a tetracycline efflux pump in staphylococci.

**FIG 6.**
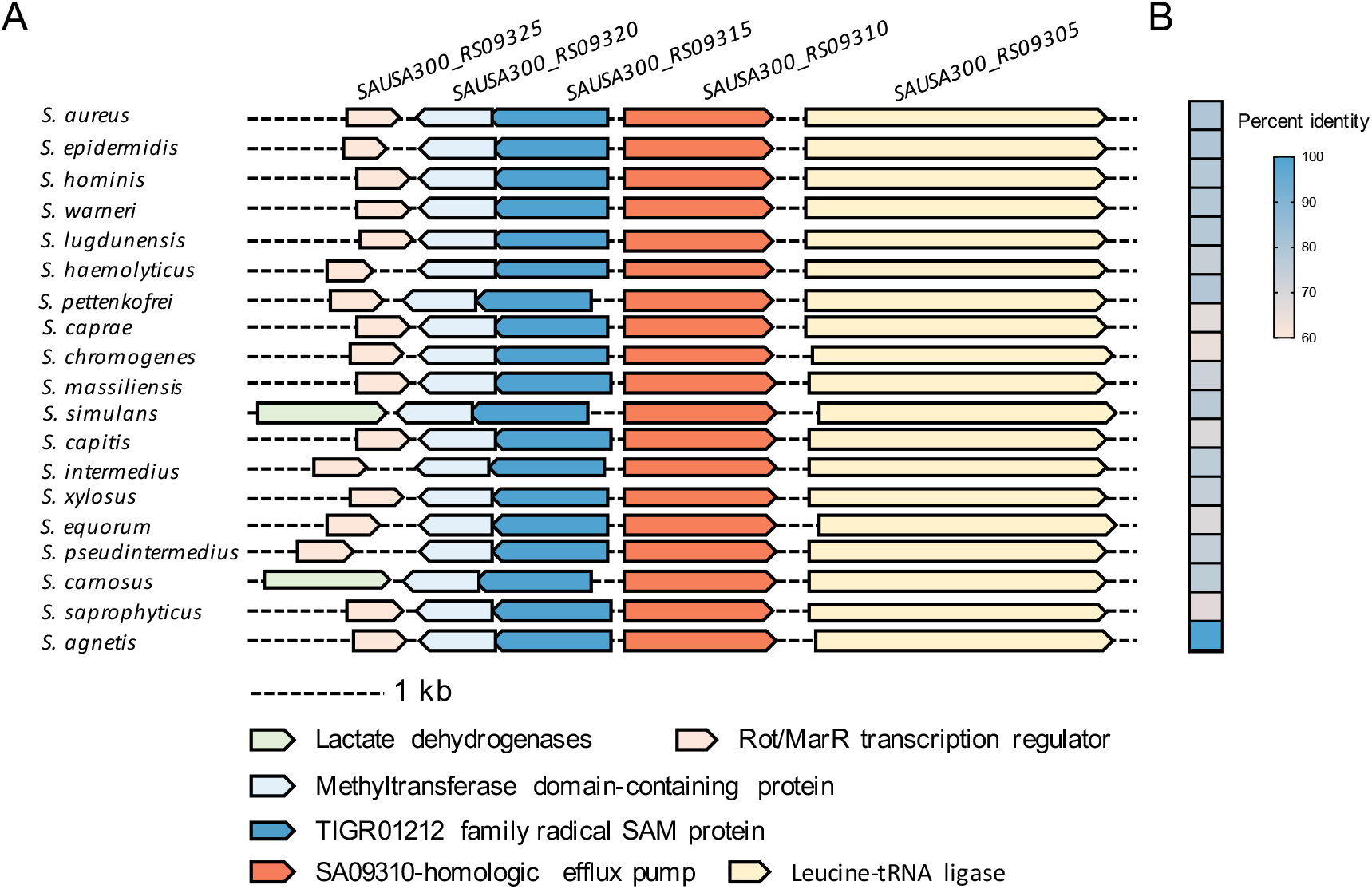
Conservation of SA09310 efflux pump in staphylococci. (A) Conserved genomic context of *sa09310* and its homologous genes in different Staphylococcus species. (B) Heatmap of percentage identities between the SA09310 protein and its analogous proteins from other Staphylococcus species.

## Materials and methods

### Bioinformatics analysis

The transmembrane helices of the SA09310 protein were predicted by using the TMHMM 2.0 online tool (http://www.cbs.dtu.dk/services/TMHMM/). The transmembrane topology was generated by TOPO2 (http://www.sacs.ucsf.edu/cgi-bin/open-topo2.py/) based on the predicted transmembrane structure from TMHMM 2.0. The genomic context of *sa09310* and its homologous genes in different Staphylococcus species was generated from the STRING database (https://string-db.org/).

To generate the phylogenetic tree of SA09310 within the MFS transporter family, at least two or more protein sequences from each of the 14 classified groups of MFS transporters, including nitrate-nitrite porter (NNP), drug-H^+^ antiporter with 14-spanner efflux (DHA14), phosphate-H^+^ symporter (PHS), nucleoside-H^+^ symporter (NHS), sialate-H^+^ symporter (SHS), metabolite-H^+^ symporter (MHS), oligosaccharide-H^+^ symporter (OHS), oxalate-formate antiporter (OFA), anion-cation symporter (ACS), oxalate-formate antiporter (OFA), organophosphate-inorganic phosphate antiporter (OPA), drug-H^+^ antiporter with 12-spanner efflux (DHA12), monocarboxylate porter (MCP), fucose-galactose-glucose-H^+^ symporter (FGHS), and unknown major facilitator (UMF) (17), were downloaded from UniProt. Multiple sequence alignments between SA09310 and MFS transporters were performed by using ClustalW (24), and the phylogenetic tree was generated from multiple alignments using the neighbor-joining method and Jukes-Cantor protein distance model, as implemented in CLC Main Workbench (Qiagen).

### Strains and growth conditions

The *S. aureus* USA300_FPR3757 strain was used to generate all the *S. aureus* mutants. Plasmids used for *S. aureus* USA300 transformation were modified by *S. aureus* RN4220. All *S. aureus* transformants were obtained through electroporation as described previously (25). *S. aureus* strains were cultured in tryptic soy broth (TSB) with shaking at 220 rpm or on TSB agar plates (TSA) at 37°C. DH5α was used for plasmid cloning and was grown in Luria-Bertani (LB) broth with constant shaking at 220 rpm or on LB agar plates at 37°C. Antibiotics were added where indicated at the following concentrations: 100 µg/mL ampicillin, 10 µg/mL tetracycline, and 25 µg/mL chloramphenicol.

### DNA manipulations

Genomic isolation and plasmid preparation from *E. coli* or *S. aureus* were performed as previously described (26). PCRs were performed using OneTaq 2× Master Mix or Q5 High-Fidelity 2× Master Mix from NEB according to the manufacturer’s instructions. Cloning was performed by using the ClonExpress II One Step Cloning Kit (Vazyme, Nanjing, China) based on homologous recombination. All primers used in this study are listed in Supplementary Table 1.

### Generation of *sa09310* knockout, complementary, and overexpression strains

Knockout of the *sa09310* gene was achieved by homologous recombination. DNA fragments of 1 kb flanking the *sa09310* gene were amplified by PCR using the primer pairs QL1165/QL1166 and QL1167/QL1168. These two DNA fragments were fused by fusion PCR via their overlap sequence. The fused fragment was cloned into the *S. aureus*-*E. coli* shuttle vector pBT2 (27) linearized with EcoRI and SalI restriction enzymes. The generated plasmid was modified by RN4220 and subsequently transformed into *S. aureus* USA300 WT with the selection of chloramphenicol. Since pBT2 carries a temperature-sensitive replicon, the plasmid was forced to integrate into the chromosomal DNA upstream or downstream of the *sa09310* gene by switching the temperature from 37°C to 42°C with chloramphenicol selection. After the correct genotype was confirmed by PCR using primers QL1169/QL1170, the resulting strain was cultured in TSB at 25°C to promote the second round of homologous recombination without antibiotics. The double crossovers were counter-selected on the basis of chloramphenicol sensitivity. The *sa09310* gene deletion mutant (Δ*sa09310*) was confirmed by PCR and sequencing. The complementary strain of Δ*sa09310* (Δ*sa09310_*com) was generated by complementing the *sa09310* gene at its original site on the chromosome in Δ*sa09310* using the same method as gene knockout. Overexpression of *sa09310* was achieved by using the replicative vector pQLV1025 with the strong constitutive promoter P_*08825*_ from *S. aureus* (26).

### Antibiotic stimulation and RT-qPCR

Cells of *S. aureus* from the mid-exponential phase were stimulated with 100 µg/mL of different antibiotics, which included quinolone (norfloxacin), glycopeptide (vancomycin), beta-lactam (ampicillin, oxacillin), aminoglycoside (gentamicin, kanamycin), tetracycline, macrolides (erythromycin), and chloramphenicol. After incubation for 1 hour, bacterial cells were harvested by centrifugation. Total cellular RNA was isolated by using the RNApure Bacteria Kit (CwBIO, Jiangsu, China) following the manufacturer’s instructions. Approximately 1 µg of total RNA was used for reverse transcription using the PrimeScript™ RT reagent kit with gDNA Eraser (Takara, Beijing, China). After the transcribed cDNAs were 5-fold diluted, 2 µL of the cDNA was used as DNA template in 15-µL amplification volumes with 400 nM of each primer and 7.5 µL of SYBR green master mix (Takara, Beijing, China) using the following cycling parameters: 95°C 30 s; followed by 40 cycles of 5 s at 95°C, 30 s at 55°C, and 30 s at 72°C. The qPCR was performed in a CFX-96 Touch Real-Time PCR system (Bio-Rad, Hercules, CA, United States). Primer pairs QL1293/QL1294 and QL0152/QL0153 were used to amplify the *sa09310* and *16S rRNA* genes, respectively. The expression level of the *sa09310* gene under the treatment with different antibiotics was normalized to the expression of the *16S rRNA* gene.

### Antimicrobial susceptibility assay

The disk diffusion assay was performed to test the susceptibility of *S. aureus* to a broad range of antibiotics. Bacterial cells from the mid-exponential phase were harvested and adjusted to an OD_600_ value of 1.0 in TSB liquid medium. One hundred microliters of the suspension were mixed with 10 mL of soft TSA (5% agarose) and spread on a TSA plate. The antimicrobial-impregnated disks were placed on the surface of the agar. Antibiotic disks with a 6 mm diameter were purchased from Microbial Regent Company (Hangzhou Microbial, Hangzhou, China). The type and amount of the antibiotic for each disk were listed in table 1. After incubation at 37°C for 24 h, the zones of inhibition around the disks were recorded according to previously reported guidelines (28).

For the MIC test, the MICs of tetracycline and doxycycline were determined by the E-test method. Bacteria-spreaded TSA plates were prepared as described above.

Tetracycline or doxycycline MIC test strips were purchased from Liofilchem (Abruzzi, Italy) with a gradient antibiotic concentration from 0.016 to 256 µg/mL. The strip was placed on the agar surface using forceps. After incubating the plate at 37°C for 24 h, the MIC value was read by viewing the symmetrical inhibition ellipse on the plate according to the guidelines given by the Clinical and Laboratory Standards Institute (CLSI).

### Intracellular tetracycline assay

The day culture of *S. aureus* was prepared by inoculating 100 µL of the overnight culture into 10 mL of TSB medium. After 2 h of cultivation, a final concentration of 5 µg/mL tetracycline was added to the culture. One milliliter of bacterial cells was collected after 0.5, 1, 1.5, 2, 3, 4, or 5 h of incubation with tetracycline. Cells were washed 3 times with PBS and resuspended in the lysis buffer (20 mM Tris-Cl, pH 8.0; 2 mM sodium EDTA; 1.2% Triton X-100) with the OD_600_ value adjusted to 1.0 (5×10^8^ CFU/mL). One hundred microliters of the suspension were used and incubated with a final concentration of 50 µg/mL lysostaphin at 37°C until the bacterial cells were completely lysed. Intracellular tetracycline released into the lysis was measured by using a tetracycline ELISA kit (Ruixin Biotech, Quanzhou, China) according to the manufacturer’s instructions. Briefly, the supernatant of the cell lysate was mixed with HRP-labeled anti-tetracycline antibody and then added into the well on a 96-well plate that pre-coated with tetracycline. Tetracycline from lysis or the precoated well will competitively interact with the antibody. After incubation at 37°C for 30 mins, each well was washed with 100 µL PBS 5 times. Next, 100 µL of substrate solution was added to each well and incubated at 37°C for 15 mins. The reaction was stopped with 50 µL stop solution, and the absorbance value of each well at 450 nm was read and recorded. Absorbance values and concentrations of tetracycline standard samples were used to generate the four-parameter logistic standard curve. The concentration of tetracycline from cell lysis was calculated according to the absorbance value based on the equation of the standard curve.

### Ethidium bromide (EtBr) accumulation and efflux assays

For the EtBr accumulation assay, *S. aureus* cells from the mid-exponential phase were harvested through centrifugation at 10,000 g for 5 min, washed with PBS 3 times, and resuspended at an optical density at OD_600_ = 0.4 in PBS supplied with 10 mM glucose. One hundred microliters of the suspension were added in triplicate to a 96-well black plate with clear bottom for measurement of the baseline cellular fluorescence for 2 mins. Following this, EtBr was added to each well to a final concentration of 4 µg/mL, and fluorescence was measured every 30 s using a Synergy H1 plate reader (Bio-Tek) at emission and excitation wavelengths of 580 nm and 500 nm respectively. For the EtBr efflux assay, *S. aureus* cells were treated with EtBr as described above for 40 mins. Then, the extracellular EtBr was removed by centrifugation and resuspension of the cells in fresh PBS, and the fluorescence of each strain was measured in a 96-well black plate by the Bioreader. After 20 mins, the efflux pump inhibitor carbonyl cyanide-chlorophenylhydrazone (CCCP) at a final concentration of 100 µM was added to each well, and the fluorescence was monitored for additional 40 mins.

### Clearance assay of *S. aureus* by tetracycline in *G. mellonella* larvae infection model

Bacterial inocula were prepared by diluting 100 µL overnight cultures with 10 mL fresh TSB followed by incubation for 2 h on an orbital shaker at 37°C to obtain bacteria in the exponential growth phase. Bacterial cells were harvested by centrifugation and resuspended in PBS at an optical density of OD_600_ = 1.0 (5 × 10^8^ CFU/mL). *G. mellonella* larvae were originally obtained from JingmaiBio (JingmaiBio, Chengdu, China), further bred in our laboratory, and used at a weight between 350 and 400 mg. Twenty larvae in each group were then inoculated with 10 µL of different amounts of bacterial suspensions (10^4^, 10^5^, 10^6^, or 10^7^ CFU/larva) into the last right proleg using a 25 µL Hamilton syringe (Sangon, Shanghai, China). After injection, larvae were incubated at 37°C for 4 days, and survival was recorded daily.

For the *S. aureus* clearance assay, 40 larvae in each group were infected with the optimal infection dose of each strain as described above. A total amount of 10 µg tetracycline was administered 10 µL into the last left proleg within 2 h after infection. Treatment was given only once, and PBS was administered as a control group for antibiotic treatment. Larvae were incubated in Petri dishes at 37°C. Five live larvae were randomly selected from each group and were tested for bacterial burden at 12 and 24 h after infection. Briefly, the larva was externally disinfected with 75% ethanol, dried, and then placed into a 5 mL tube with 2 mL sterilized PBS. Larvae in the tube were completely homogenized by using a portable homogenizer with a 4 mm tip (PRIMASCI, UK). Larval homogenate (50 µL) was serially diluted in 450 µL of PBS. The dilution series was plated on TSA agar plates and was incubated at 37°C for 24 h. Colonies were counted after 24 h, and data are expressed as CFU per larva.

## Discussion

In this study, we investigated the role of the *sa09310* gene in mediating antimicrobial resistance of *S. aureus* USA300. The *sa09310* gene was predicted to encode an MFS transporter with 12 TMS, which exhibited a close relationship to the multidrug efflux pump DHA12 according to the phylogenetic analysis. Deletion of the *sa09310* gene renders *S. aureus* more sensitive to tetracycline both *in vitro* and *in vivo*. In the presence of tetracycline, the Δ*sa09310* mutant exhibited a higher concentration of intracellular tetracycline than the WT. Additionally, an EtBr accumulation assay was performed to prove that SA09310 displayed efflux pump activity. All this evidence indicated that SA09310 was a tetracycline efflux pump that mediated tetracycline resistance in *S. aureus*.

Currently, the multidrug efflux pumps are classified into five families, among which the MFS has been the most extensively studied among staphylococci, which include NorA, NorB, NorC, Tet38, LmrS, SdrM, and MdeA (14). The characteristics of MFS transporters are their possession of 12 or 14 TMS (17)., and two highly conserved motifs, called motif A and motif C, consisting of residues “G(X)_3_DK/RXGRR/K” and “G(X)_8_G(X)_3_GP(X)_2_GG”, which lie in the cytoplasmic loop between the TMS2 and TMS3 helices and the fifth α-helix (TMS5) (Fig. 1A). The same signatures of MFS were found in SA09310 protein (Fig. 1A). All this evidence strongly suggested that the *sa09310* gene encoded an MFS transporter. MFS transporters so far have been classified into 17 distinct groups based on their sequence similarity and their functional relevance (17). To predict the function of SA09310 transporter, a phylogenetic tree was constructed based on the protein sequence alignment between SA09310 and the classified MFS transporters. SA09310 exhibited a close relationship to the drug-H^+^ antiporter (12-spanner) drug efflux (DHA12) and monocarboxylate porter (MCP) families (Fig. 1A), which indicated that SA09310 was probably involved in drug or carboxylate efflux.

The expression of most drug efflux pump-encoding genes, such as *qarC* and *abcA* in *S. aureus*, can be induced by their target substrate (29, 30). To identify the substrate transported by SA09310, we first checked the response of the *sa09310* gene to the stimulus of different types of antibiotics. Surprisingly, the expression of *sa09310* was induced by most of the tested antibiotics (Fig. 2A). This observation implied that the SA09310 efflux pump might be involved in the extrusion and resistance to these antibiotics without specificity. Unexpectedly, the susceptibilities of *S. aureus* WT and Δ*sa09310* mutant displayed no difference to most of these antibiotics except tetracycline and doxycycline. It seems that there is no necessary association between antibiotic induction and efflux pump-mediated corresponding resistance.

The multidrug efflux systems in bacteria are often regulated by transcription factors that themselves bind the substrates of these export systems thereby allowing them to activate or repress the expression of their cognate transporter genes (31). One significant family of such transcription factors is the multiple antibiotic resistance repressor (MarR) family. MarR coding genes are often located in the nearby gene of its controlled transporter-encoding genes, such as the typical genes MarA and MprA in *E. coli* (32, 33), MexR in *P. aeruginosa* (34), and MepR in *S. aureus* (35).

Although most of the multidrug efflux genes were located next to MarR coding genes, there is no such a *marR* gene nearby *sa09310* gene. Instead, a putative MarR coding gene with the gene_lucos tag SAUSA300_RS09325 that was separated by two genes upstream of *sa09310* was identified (Fig. 6A). Whether the expression of the *sa09310* gene is regulated by this MarR regulator, and the regulatory mechanism of tetracycline-induced expression need further investigation.

Tetracyclines are considered as antimicrobial agents since they preferentially bind to intracellular ribosomes and interact with a highly conserved 16S ribosomal RNA (rRNA) target in the 30S ribosomal subunit, arresting translation by sterically interfering with the docking of aminoacyl-transfer RNA (tRNA) during elongation (21-23). Tetracycline resistance in bacteria is currently attributed to three well-known mechanisms: active efflux, ribosomal protection, and enzymatic inactivation of tetracycline (36). To date, transporters NorB and Tet38 from the MFS family and MepA from the MATE family have been reported to be involved in tetracycline antibiotic resistance in *S. aureus* (35, 37, 38). The mechanism of these transporters mediating resistance was attributed to their ability to export tetracycline (35, 37, 38). The Δ*sa09310* mutant displayed a significantly lower concentration of intracellular tetracycline (Fig. 4A), suggesting that the mechanism of SA09310-mediated resistance to tetracycline is similar to the mechanism that mediated by Tet38, NorB, and MepA through the extrusion of the substrate (35, 37, 38). This efflux activity of SA09310 was further verified by EtBr accumulation and efflux assays. Although SA09310 promoted *S. aureus* resistance to tetracycline and doxycycline, the Δ*sa09310* mutant did not show increased susceptibility to tigecycline, which belonged to tetracycline antibiotics (data not shown). This observation implied that SA09310 recognized and extruded its substrate with high specificity even in the same type of antibiotic.

Knockout of the *sa09310* gene conferred *S. aureus* with a 64-fold decreased tetracycline MIC *in vitro*, which implied that the SA09310 efflux pump might be utilized as a potential target of an antibiotic adjuvant of tetracycline thereby to overcome efflux-based resistance. However, the increased susceptibility of the Δ*sa09310* mutant to tetracycline *in vitro* might not necessarily predict *in vivo* outcomes. Bearing that in mind, we investigated the clear efficiency of tetracycline against *S. aureus* in a *G. mellonella* larvae infection model *in vivo*. As expected, the Δ*sa09310* mutant was cleared by tetracycline more easily compared to WT in *G. mellonella* infection model. This provided compelling evidence for the feasibility of developing and using a SA09310-targeted inhibitor as an antibiotic adjuvant to overcome efflux-based tetracycline antibiotic resistance in the clinic.

Taken together, we identified a novel and staphylococci-conserved MFS transporter and demonstrated its role as a tetracycline efflux pump that mediates tetracycline resistance in *S. aureus*. Further studies should be carried out to decode the regulatory mechanism of *sa09310* expression.

## Acknowledgments

This work was supported by the National Science Fund for Distinguished Young Scholars (32000094), Sichuan Natural Science Fund for Distinguished Young Scholars (2022NSFSC1682), the China Postdoctoral Science Foundation (2021M692311), the Post-doctor Research Project, West China Hospital, Sichuan University (20HXBH017), and the 1·3·5 project for disciplines of excellence, West China Hospital, Sichuan University (ZYXY21004).

